# Evolution of protein kinase substrate recognition at the active site

**DOI:** 10.1101/443945

**Authors:** David Bradley, Pedro Beltrao

## Abstract

Protein kinases catalyse the phosphorylation of target proteins, controlling most cellular processes. The specificity of serine/threonine kinases is partly determined by interactions with a few residues near the phospho-acceptor residue, forming the so-called kinase substrate motif. Kinases have been extensively duplicated throughout evolution but little is known about when in time new target motifs have arisen. Here we show that sequence variation occurring early in the evolution of kinases is dominated by changes in specificity determining residues. We then analysed kinase specificity models, based on known target sites, observing that specificity has remained mostly unchanged for recent kinase duplications. Finally, analysis of phosphorylation data from a taxonomically broad set of 48 eukaryotic species indicates that most phosphorylation motifs are broadly distributed in eukaryotes but not present in prokaryotes. Overall, our results suggest that the set of eukaryotes kinase motifs present today was acquired soon after the eukaryotic last common ancestor and that early expansions of the protein kinase fold rapidly explored the space of possible target motifs.

## Introduction

Protein kinases are essential for signal transduction and have been found in every eukaryotic species so far examined. They are required for almost all cellular processes (Endicott, Noble, and Johnson 2012), and mutations in protein kinases are often associated with diseases such as cancer and diabetes (Stenberg, Riikonen, and Vihinen 2000; Lahiry et al. 2010; Torkamani et al. 2008). Kinases are often described in terms of their ‘specificity), which refers to the set of substrates that the kinase is able to phosphorylate *in vivo.* Multiple factors define the specificity of the kinase (Ubersax and Ferrell 2007). The kinase and substrate must be co-expressed and co-localised for example, and their interaction may be mediated by adaptor or scaffold proteins (Pawson and Scott 1997; Faux and Scott 1996). Docking sites on the substrate may also be employed to recruit the kinase and the substrate directly (Biondi and Nebreda 2003; Goldsmith et al. 2007). Fundamentally, selectivity is often defined by the structural interface between the kinase active site and the residues flanking the target serine, threonine, or tyrosine – the so-called ‘peptide specificity) of the kinase.

The kinase peptide specificity is usually described in terms of a short linear motif (Pearson and Kemp 1991; Pinna and Ruzzene 1996). The substrate motif of PKA, for example, is R-R-x-S/T, meaning that an arginine is preferred 2 and 3 positions N-terminal to the target serine/threonine in the PKA active site. Conceptually, different substrate motifs can be thought of as different channels of communication within the cell, allowing for kinases that are simultaneously active to regulate a specific set of substrates. Mitotic kinases with overlapping localisations, for example, tend to have mutually exclusive substrate motifs, presumably to prevent the aberrant phosphorylation of non-targets during cell cycle progression (Alexander et al. 2011).

The range of possible selectivities at the active site is likely restricted by the structure of the kinase domain itself. In turn, the capacity of the kinase fold to create novel specificity preferences through mutations may determine the maximum effective number of kinases possible for a genome, as it has been suggested for transcription factors (Itzkovitz, Tlusty, and Alon 2006). In this analogy to a communication channel, the full set of possible substrate motifs can be thought as the full bandwidth or “communication potential” of the kinase fold. Understanding how this communication potential was explored over evolutionary time can reveal insights into the evolution of cell pathways and cell signalling. However, while the proliferation of the kinase domain itself has been well documented, much less is known about the evolution of new kinase specificities at the active site (Ochoa, Bradley, and Beltrao 2018). One study found that the frequency of tyrosine kinases in the proteome correlates negatively with the frequency of tyrosine residues in the proteome, implying some extent of coevolution between kinases and substrates (Tan et al. 2009). Another study found that the evolution of a new specificity in the CMGC group proceeded through an intermediate of broad specificity (P+1/R+1) before later specialisation into distinct target preferences (P+1 and R+1) (Howard et al. 2014).

Currently, the scarcity of kinase-substrate interaction data outside of a few model organisms (human, mouse, and budding yeast) is limiting for further research. However, other sources of data can yield insights more indirectly. An evolutionary analysis of the kinase domain can be informative provided that the specificity-determining positions (SDPs) are known (Bradley et al. 2017). This applies to phosphoproteome data also provided that motifs can be extracted and linked to the known specificities of kinase families or subfamilies. Here we collect kinase sequence data, kinase specificity data, and phosphorylation data from several species to perform an evolutionary analysis of kinase specificity. Collectively, the results suggest that most specificities arose early in the evolution of protein kinases, followed by a long period of relative stasis.

## Results

### Residues implicated in the differentiation of duplicated kinases

The eukaryotic protein kinase superfamily by convention is divided hierarchically at the level of ‘groups), ‘families), and ‘subfamilies) (Hanks and Hunter 1995; Manning et al. 2002). The eight canonical kinase groups (AGC, CAMK, CK1, CMGC, RGC, STE, TKL, TK) evolved the earliest and, with the exception of tyrosine kinases (TKs) and the RGC group, are thought to have arisen in an early eukaryotic ancestor (Miranda-Saavedra and Barton 2007). Kinase families and then subfamilies generally emerged later during evolution and reflect more distinct features of the kinase)s function (specificity, regulation, localisation, *etc*) (Hanks and Hunter 1995). In order to study the evolution of kinase specificity, we first performed a systematic phylogenetic analysis to predict kinase functionally divergent residues for every kinase family and subfamily where possible. To this end, a global kinase domain phylogeny was constructed for the 9 annotated opisthokont kinomes (*H. sapiens*, *M. musculus*, *S. purpuratus, D. melanogaster, C. elegans, A. queenslandica, M. brevicollis, S. cerevisiae,* and *C. cinerea*) present in the kinase database *KinBase* **(http://kinase.com/web/current/kinbase/).** For 99 kinases families and 88 kinase subfamilies, we identified residues that are conserved within a clade but differ from the sister clade in the phylogeny. These residues are implicated as functionally divergent residues and are expected to underlie functional differences between kinase sister clades. This was achieved by calculating divergence scores (*s)* for each alignment position and each family and subfamily using an adaptation of the BADASP method (Edwards and Shields 2005) that is based on ancestral sequence reconstructions (**Figure 1a, Methods**).

**Figure 1.**
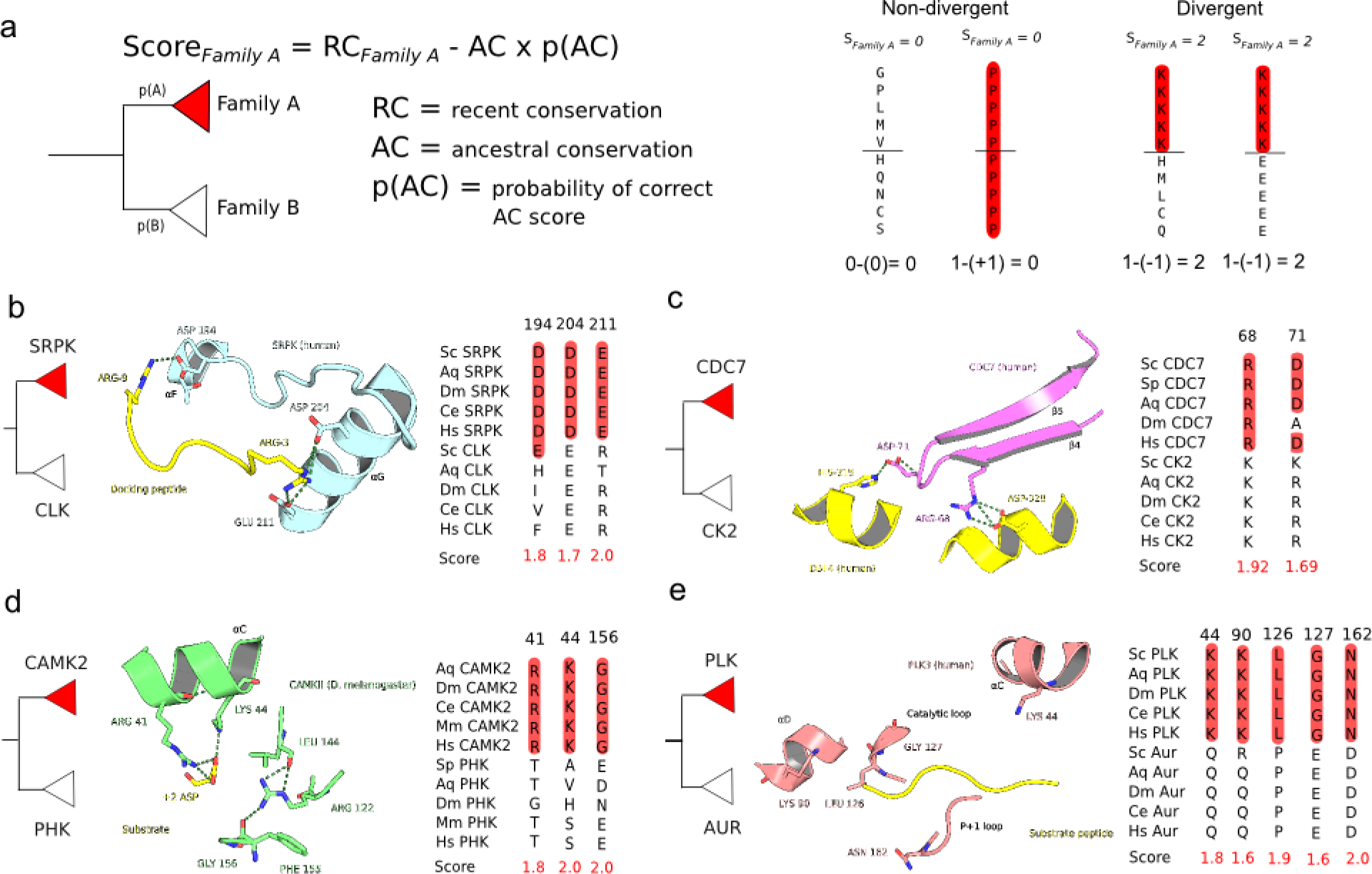
Family and subfamily divergent residues. a) Explanation of the score used to identify divergent residues. ***RC (recent conservation):*** conservation of residue within the family of interest. ***AC (ancestral conservation):*** extent to which the residues are conserved between sister clades in the phylogeny. ***p(AC):*** a measure of confidence in the ancestral sequence prediction. A full explanation is given in the ***Methods*** section. ***b-e)*** Examples of residues predicted to be functionally divergent in the SRPK, CDC7, CAMK2, and PLK families.

Functionally divergent residues were first predicted across all kinase families. A detailed analysis of the results suggests multiple ways in which novel kinase functions have evolved at the family level via changes in functionally relevant residues. In the SRPK family (CMGC) for example, two substitutions to negatively charged amino acids (D and/or E) map to parts of the kinase structure that have been shown previously to bind to a positively charged docking peptide (**Figure 1b**; (Ngo et al. 2005)). In the CDC7 family (CMGC) also, two of the identified functionally divergent residues bind to the CDC7 activator protein named Dbf4 (**Figure 1c**; (Hughes et al. 2012)), and are therefore important for kinase regulation. In other examples, the functionally divergent residues identified can help to account for the specificity of the kinase. Two substitutions in the CAMK2 family (CAMK) for example bind to a preferred D/E residue at the substrate +2 position **(PDB: 5H9B, unpublished)**. A glycine substitution in the activation loop has also been shown to be important for kinase function (LeBoeuf, Gruninger, and Garcia 2007), and may explain why CAMK2 kinases do not require activation loop phosphorylation for activity ((Bhattacharyya et al. 2016), **Figure 1d**). Finally, many substitutions for the acidophilic PLK family map to SDRs and are convergent with those identified for the unrelated GRK family (AGC) that is also acidophilic ((Bradley et al. 2017), **Figure 1e**). These examples illustrate how the predicted functionally divergent sites between families or subfamilies of kinases can map to functionally relevant residues. We next studied if this would be a general feature of these residues across many families and subfamilies.

### Functionally important residues are often divergent across kinase families and subfamilies

Across all kinase families, we aggregated the total number of predicted functionally divergent residues or ‘switches’ at each position in the kinase domain. This allows us to predict positions that often determine the functional differences between kinase families. These were mapped across the kinase catalytic domain sequence and fold (**Figure 2, left**). The distribution of residues that are often implicated in kinase family functional differences is not uniform and strongly enriched within or close to the kinase activation segment, the αC helix, the β5-αD region, and the αF-αG regions (**Figure 2, left**). For further analysis, we divided kinase residues into functional categories: ‘catalytic) (catalytic residues and the catalytic spine), ‘proximal) (within 4 Angstroms of the peptide substrate), ‘distal SDRs) (distal SDRs implicated in (Bradley et al. 2017)), ‘regulatory) (regulatory spine residues and those within and surrounding the activation loop), ‘interaction) residues (those most frequently in contact with other protein domains) and ‘other) (residues not belonging to any of the previous categories). We defined as frequently-switching residues those at the 90th percentile of residues with most changes. The majority (14/21) of frequently switching residues can be assigned to a functional category (catalysis, specificity, regulation, etc.), which is more than would be expected by chance (p=0.0083; Fisher)s Exact Test, one-sided). This suggests that this approach can successfully identify residues that are of relevance for the functional divergence of kinases. Of these residues, 8 have been implicated in determining differences in kinase specificity. The number of substitutions for specificity determining residues is generally higher than that for residues without an assigned function (Mann-Whitney, one-tailed, p = 1.1×10^-5^). These results suggest that substrate-determining residues often undergo substitutions as new kinase families emerge.

**Figure 2.**
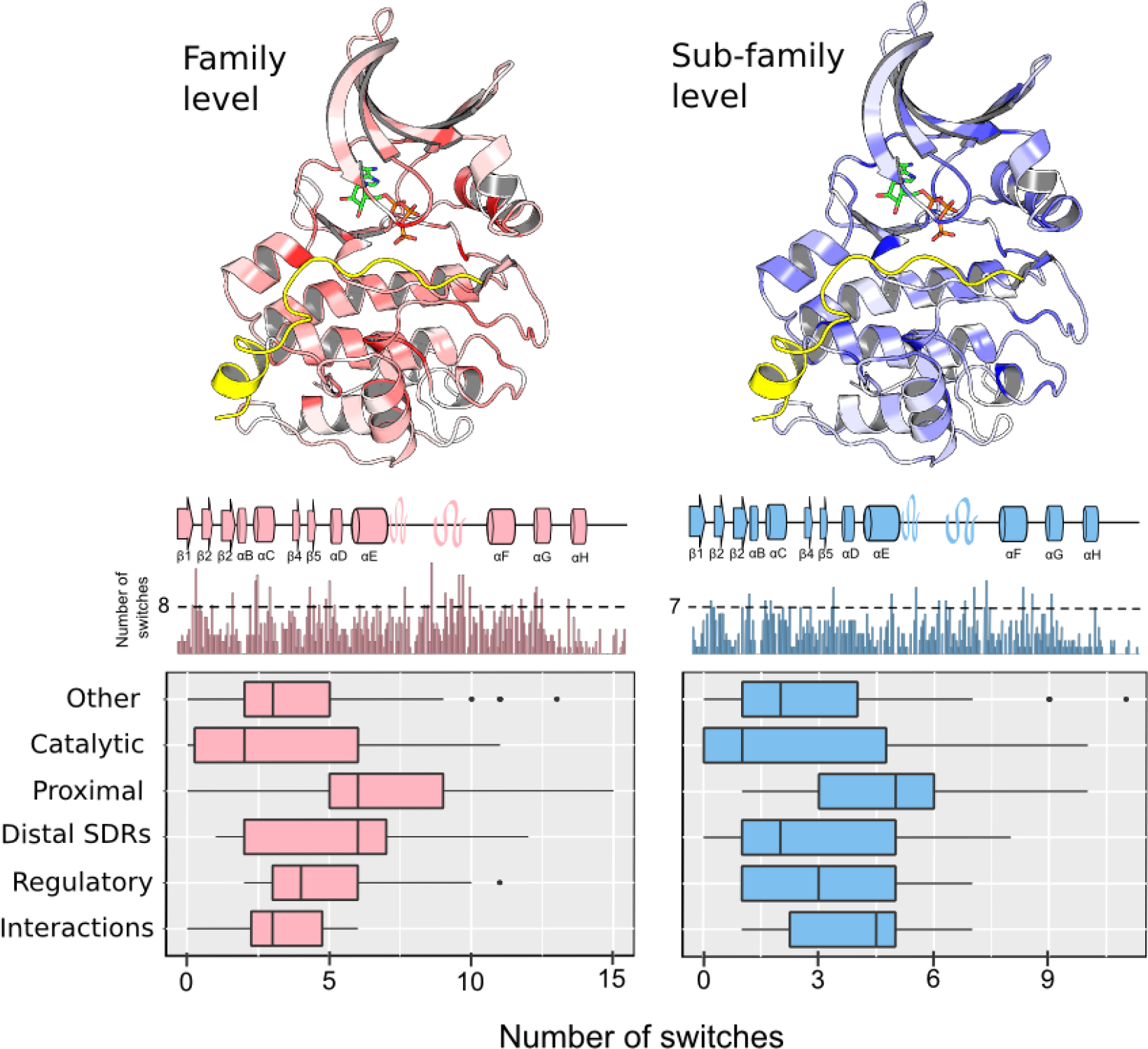
Aggregated analysis of sequence divergence across kinase families (left) and subfamilies (right). For each kinase domain position, the total number of ‘switches’ (score > 95th percentile of scores) was counted across all families or subfamilies considered (see ***Methods***). ***Top:*** Results mapped to kinase structures. Darker shades of red/blue represent a higher number of switches. ***Middle:*** Total number of switches mapped to the kinase primary sequence, with secondary structure elements represented above the barplot. A domain position is considered to be ‘frequently switching’ if the number of switches lies above a 90th percentile threshold for the kinase domain (246 positions). The threshold is ‘8’ for families and ‘7’ for subfamilies. ***Bottom:*** The values for each domain position have been grouped according to the functional category (‘catalytic’, ‘regulatory’, ‘proximal’, etc) and the distribution plotted separately at the family and subfamily level.

A similar analysis was performed for kinase subfamily comparisons (**Figure 2, right**). Similar to kinase family evolution, a large fraction (73%, 11 out of 15) of residues frequently implicated in the functional differences between kinase subfamilies were also annotated to a functional category (p=0.0068; Fisher)s Exact Test, one-sided). We also observed a higher than expected number of switches for putative specificity determining residues at the subfamily level when compared to residues not annotated with a function (Mann-Whitney, one-tailed, p = 1.0×10^-4^). However, we note that the frequently switching residues for subfamilies are more evenly distributed across functional classes than for the family level analysis.

### Evolution of experimentally determined kinase target preferences

The above results suggest that residues important for kinase specificity are often different across kinase families and less so across subfamilies. We then studied the extent by which these changes in kinase residues impact on their target specificity. To study this, we derived kinase specificity models for 101 S/T kinases from human and mouse using experimentally determined target sites (**Methods**). We then tested the extent to which kinase specificities differ within and between groups, families, and subfamilies for kinases of known specificity. In line with expectation, the differences in specificity are larger across groups than across families, and also larger across families than across subfamilies (**Figure 3a).** For subfamilies, the differences are not statistically different from the distances measured within subfamilies (p=0.21, Kolmogorov-Smirnov-test, two-sided). These results suggest that kinase specificity often diverges at the level of the group, less so at the family level, and rarely when new subfamilies emerge. We show in **Figure 3b** some examples of typical differences in kinase specificity for the 3 classifications. Although the differences in specificity across families is statistically different from expectation (p<<0.01, Kolmogorov-Smirnov-test, two-sided), the “typical” differences observed are smaller than at the group level. This is illustrated by the RSK and PKC families **(Figure 3b, center),** which both have a preference for arginine at the -3 position, but PKC additionally has a preference for R at position +2 while RSK has a modest preference for the same residue at position -5.

**Figure 3.**
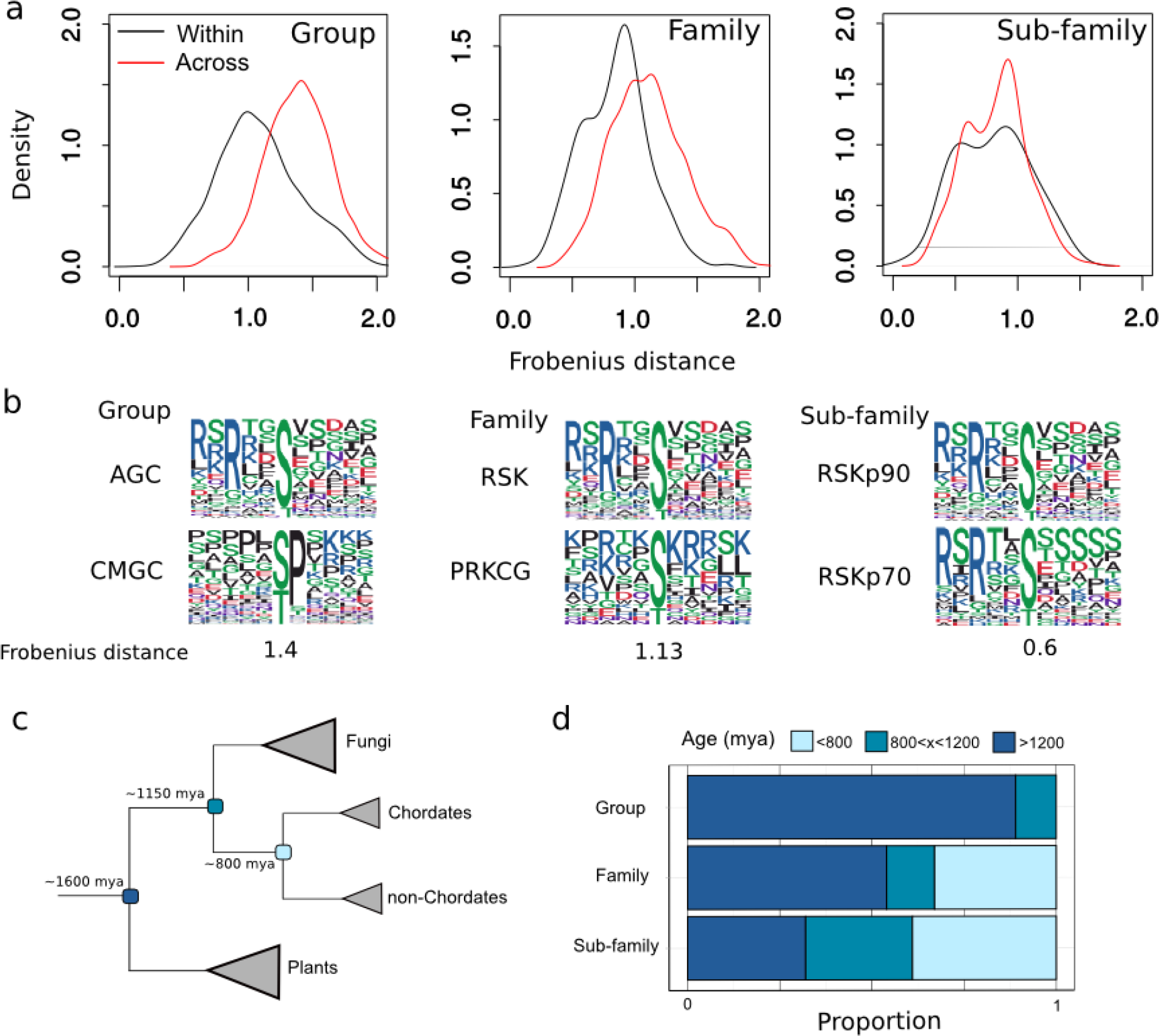
**a)** Differences in S/T kinase specificity models at the group, family and subfamily levels. The Frobenius distance was calculated for all possible pairwise comparisons within and between groups, families, and subfamilies ***b)*** Representative kinase pairs belonging to different groups (left), families (center), and subfamilies (right). Frobenius distances for each of the 3 pairs are given beneath the logos ***c)*** A simplified Tree of Life with three important divergence times (plant-opisthokont, fungi-metazoa, chordate - non-chordate) marked. ***d)*** Phylogenetic estimation of kinase ages at the group, family, and subfamily level for S/T kinases.

To put the previous results into the context of evolutionary time-scales, we sought to estimate the time of origin for many kinase groups, families and subfamilies. To this end, the presence or absence of every S/T kinase group, family, and subfamily, was predicted for several species across the Tree of Life **(Figure 3c)**. The phylogenetic origin of kinase groups/families/subfamilies was then predicted using ancestral state reconstruction, which allowed their emergence to be dated based on the know divergence times between species (Kumar et al. 2017). Overall, we estimated that most kinase groups arose in a universal eukaryotic ancestor, in line with a previous study (Miranda-Saavedra and Barton 2007). For kinase families, around 55% are estimated to have arisen in a universal ancestor and up to 65% have arisen before the split between chordates and non-chordates (~800 mya). Around 60% of subfamilies were similarly estimated to have arisen before the split between chordates and non-chordates **(Figure 3d)**. Together with the analysis of kinase specificity differences, this result suggests that relatively few kinase specificities are likely to have arisen in the past 800 million years of kinase evolution.

### Kinase motif enrichment across 48 eukaryotic species

The analysis of kinase specificity differences described above can only be performed for kinases with many experimentally determined targets. For most kinases, this information is not available (Hornbeck et al. 2015; Bradley et al. 2017). As an alternative way to study the evolution of kinase specificity, we analyzed MS-derived phosphorylation sites from a broad range of species. The phosphoproteome of any given species represents an ensemble of kinase activities. Many of these kinases will have preferred target site sequence motifs that are required for optimal substrate phosphorylation. The signature of several different kinases may therefore be encoded in each phosphoproteome.

For this study, we were interested in determining the extent to which different kinase motifs have been exploited during the evolution of the eukaryotes. To this end, phosphoproteome data was collected from 48 eukaryotic species, including species from the alveolates (4), amoebozoa (1), excavates (3), fungi (19), heterokonts (1), metazoa (12) and plants (8). We first measured the enrichment of three well-established substrate signatures (R-x-x-S/T, S/T-P, and D/E+3) and found them to be strongly enriched in nearly all of the 48 species **(Figure 4, top)**. This suggests that these 3 common preferences are likely to have been present very early on during the evolution of the eukaryotes. To extend this to other kinase preferences, target site **S/T** sequence motifs were extracted from each species phosphoproteome using the *motif-x* tool (Schwartz and Gygi 2005). Motifs without consistent enrichment across related species were filtered from any further analysis (**Methods**). In total, 29 motifs were (**Figure 4**) identified, which account for **~**54% of all phosphosites analysed (**Supplementary Figure 1a**).

**Figure 4.**
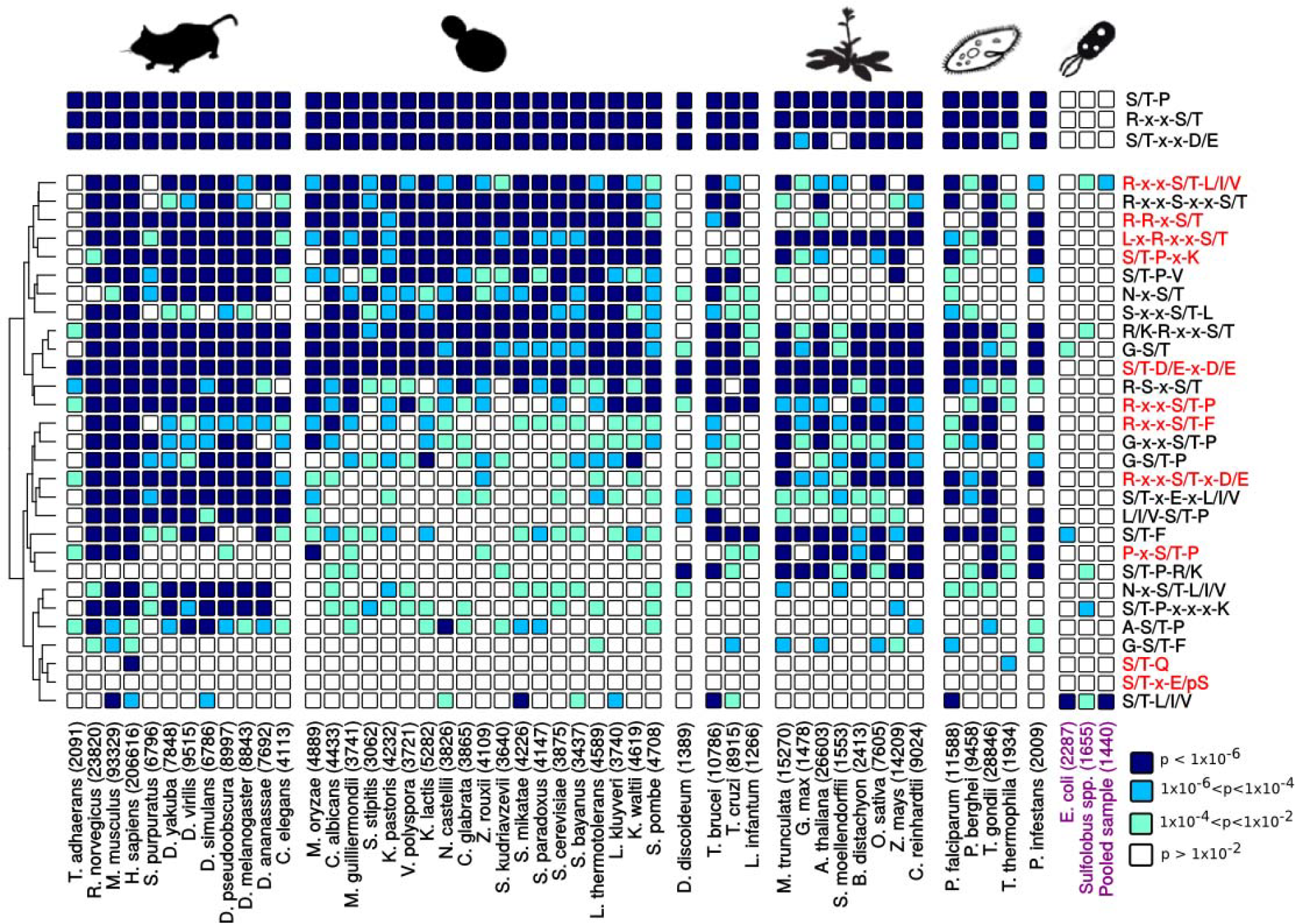
Enrichment of S/T phosphorylation motifs across several species. Binomial p-values were calculated for each motif and each species considered. The heatmap cells are coloured according to the extent of enrichment for that particular motif and species (see legend, bottom-right). The numbers in the column labels correspond to the sample size of unique S/T phosphorylation sites (15mer). Prokaryotic phosphosite samples are coloured in purple. ***Top:*** enrichment of three common phosphorylation signatures (S/T-P, R-x-x-S/T, S/T-x-x-D/E). ***Bottom:*** enrichment of 29 motifs discovered using the motif-x tool. Motifs where the effector kinase has already been described in the literature are coloured in red.

Among the 29 motifs identified, 11 have been characterised previously in the literature and have been assigned to at least one kinase family or subfamily. This includes well-known motifs such as the CDK family motif (*S/T*-P-K) and the CK2 family motif (*S/T*-D/E-x-D/E). Eight of the motifs feature either a proline at position +1 or an arginine at position -3. Other motifs were identified in addition to those that are well characterized (**Figure 4, motifs in black**). Here multiple constraints were imposed to ensure that the selected motifs were likely to represent *bona fide* kinase target motifs. For example, motifs with simple S/T additions to a classical motif were filtered from the analysis, as they could result from phosphosite misassignment within phosphopeptides from the mass spectrometry analysis or potential clustering of phosphorylation sites in the substrate primary sequences (Moses, Hériché, and Durbin 2007; Schweiger and Linial 2010). Overall, 18 uncharacterized motifs were selected using the protocol described in the *Methods* section. Some of these motifs feature ‘new’ substrate specificity determinants such as asparagine (N) and glycine (G).

In **Figure 4,** enrichment p-values for each motif were calculated for each species relative to a background set of shuffled phosphorylation site sequences, with the S/T retained at the centre. This analysis suggests that the majority of motifs (**Figure 4**) are pervasive across the eukaryotic Tree of Life. This finding is even more evident when phosphorylation data is pooled from each of the major taxonomic clades (animals, fungi, plant, etc.), and the enrichment p-values recalculated (**Supplementary figure 1b and 1c**). Most of the motifs analysed are distributed across clades that diverged early during the evolution of the eukaryotes. For example, 18 out of 29 motifs (62%) are highly enriched (p < 1×10^-6^) in animals, fungi, and plants indicating that they are therefore likely to be of ancient origin.

The distribution of motif enrichments between related species is non-random as supported by tests for the phylogenetic signal of phosphorylation motifs **(Supplementary Table 1)**. We tested whether kinase motif enrichments correlate with the frequency of their expected effector kinases in the kinome, including for example the frequency of CDKs with the frequency of S/T-P-x-K motifs. However, our analysis suggests that kinase family or subfamily frequencies are not generally correlated with motif enrichment values when the phylogenetic interdependence of data points is taken into account (Felsenstein 1985) **(Supplementary Figure 2)**. In spite of this, there is local evidence of kinase-substrate coevolution for some clades and motifs. In the plants, for example, the lack of enrichment of the basophilic R-R-x-S/T motifs can likely be explained by the depletion of its cognate effector kinase (PKA/PKG) (**Supplementary Figure 3**), as has been suggested previously (Resjö et al. 2014; Frades, Resjö, and Andreasson 2015). For the CDK family also, the pattern of S/T-P-x-K evolution and CDK evolution is similar across many species (**Supplementary Figure 4**). However, many other patterns cannot be similarly accounted for, which suggests that there are multiple factors can that affect the fold enrichment values calculated.

### Kinase motif enrichment in prokaryotes

Some kinases encoded in the genome of prokaryotes are homologous to eukaryotic protein kinases and are currently referred to as ELKs (ePK-like kinases) (Oruganty et al. 2016; Pereira, Goss, and Dworkin 2011). Current genomic data now suggests that these eukaryotic-like kinases are as prevalent as histidine kinases in the prokaryotes (Kannan et al. 2007). Until recently however the S/T phosphoproteomes of archaean and bacterial species had remained poorly characterised (M.-H.Lin, Sugiyama, and Ishihama 2015). We repeated the motif analysis on the species *E. coli* (bacteria) and *Sulfolobus spp.* (archaea), which are currently the only two organisms with more than 1,000 determined S/T phosphosites (M.-H.Lin, Sugiyama, and Ishihama 2015; Pan et al. 2015; Potel et al. 2018). This analysis suggests that a large majority of the eukaryotic phosphorylation motifs discussed previously are not significantly enriched in these two species (**Figure 4**). For *E. coli*, there is a moderate enrichment for some single-site motifs, which could have evolved convergently with those motifs found in the eukaryotes. For *Sulfolobus*, we observe only weak enrichment for five eukaryotic motifs (S/T-L/I/V, S/T-P-x-x-x-K, S/T-P-R/K, R/K-R-x-x-S/T, and R-x-x-S/T-L/I/V). We note also that the S/T-P and R-x-x-S/T motifs that are highly prevalent in the eukaryotes show no evidence of enrichment across the prokaryotic species tested (**Figure 4**). While it could be argued that the low phosphosite sample sizes (2287 and 1655 for *E.coli* and *Sulfolobus*, respectively) precludes the reliable detection of weaker motifs, we note that both of these signatures (S/T-P and R-x-x-S/T) were found to be strongly enriched in eukaryotic species with small sample sizes (*Dictyostelium discoideum*: 1389, *Leishmania infantum*: 1266). A similar lack of enrichment was found for a pooled sample of prokaryotic phosphorylation sites (n=1440) from several species (Pan et al. 2015) (**Figure 4**).

Prokaryotic motifs were then identified *de novo* using the *motif-x* tool for *E. coli* and the *Sulfolobus* genus. None of the motifs identified overlap with the motifs recovered previously for eukaryotic species (**Supplementary Tables 2 and 3**). For *Sulfolobus*, in particular, we also found that 5 out of the 7 motifs identified contain a positively charged residue (R/K), implying the existence of basophilic *Sulfolobus* kinases.

## Discussion

Here we have explored the evolution of protein kinase specificity at the active site, using a combination of kinase sequence data, phosphorylation data, and kinase specificity models. Using the sequence of protein kinases across several species, we have shown that the evolution of new kinase families is dominated by sequence changes that are likely to impact on kinase function, including kinase peptide specificity. This is in line with our observation that kinases belonging to different groups and families typically show significant differences in their specificity. In contrast, kinases belonging to sister subfamilies do not show significant differences in their specificity. A phylogenetic analysis revealed that most kinase groups and families (89% and 54%, respectively) are of ancient origin among the eukaryotes, while subfamilies generally emerged later during evolution (only 32% are of ancient origin). Finally, phosphorylation motifs determined for 48 eukaryotic species were found to be broadly distributed across divergent species and likely emerged in an early eukaryotic ancestor after their divergence from the prokaryotes. Taken these different observations together, we suggest that the majority of the kinase active site specificities present today in eukaryotic species have emerged early on during the evolution of eukaryotes.

The analysis here of divergent residues across kinase families follows a similar analysis employing a BLAST-based approach (Kalaivani, Reema, and Srinivasan 2018). This analysis is focused here on the kinase catalytic domain and we did not take into account the evolution of domain composition or sequence changes outside the catalytic domain, which may have a significant impact on catalytic function (Pearce, Komander, and Alessi 2010). Kinase docking interfaces also are not amenable to the aggregated analysis attempted here as their location in the kinase domain tends to differ significantly between families (Biondi and Nebreda 2003). In general, the approach used here assumes that a given kinase domain position will adopt a given function (catalytic, regulatory, proximal, etc) across all kinase families. Examples are known already however of modes of regulation or specificity that are particular to a given family or subfamily (Sang et al. 2018; Simon et al. 2016). The important residues may be functionally misannotated in such cases, which would underestimate the extent of divergence in regulatory or substrate-specific functions. Such kinase-specific examples of residue function may account for many of the switching events currently placed in the ‘Other’ category.

From the analysis of kinase specificity models, it is apparent that new specificities are often generated following the emergence of a new kinase group or family, but not following the emergence of a new kinase subfamily. This is not a surprising result given that kinase groups and families tend to be older than kinase subfamilies (**Figure 3d**). It is however in conflict with the finding that SDR substitutions are also present throughout the evolution of subfamilies (**Figure 2, subfamilies**). There are two known cases in the literature (PLK and GRK) where modest differences in specificity are observed between sister subfamilies (Onorato et al. 1991; Franchin et al. 2014). We suggest that differences in peptide specificity can exist between subfamilies but on average, and based on the current small sample size of specificity models, these tend to be very modest at the subfamily level.

Finally, the analysis of phosphorylation motifs across 48 different eukaryotic species suggests that most arose in an early eukaryotic ancestor. The phosphorylation dataset spans the Tree of Life but is unsurprisingly biased towards animal, fungal, and plant species. Ongoing projects for increased representation of the protist superkingdoms could help to address this problem in the future (Waller et al. 2018). From this analysis also we conclude that most eukaryotic phosphomotifs post-date the divergence of eukaryotes and prokaryotes. The acquisition of phosphoproteome data from several more prokaryote species will be required however to strengthen this conclusion. In general, the increase in statistical power enabled by larger datasets will enable the reliable identification of weakly enriched motifs (‘false negatives’), many of which are likely missing from this analysis.

Collectively, the results suggest that the evolution of new kinase specificities was characterised by a ‘burst’ in early eukaryotic evolution followed by a period of relative stasis. Most gene duplicates will be quickly silenced (Lynch and Conery 2000) and diversification of function is often considered a primary means for the “survival” a newly duplicated genes in the genome. The capacity of the kinase fold to generate diverse target preferences in the active site interaction through mutations may have been an important factor underlying the success of this fold. Our analysis suggests that over the past 800 million years there have been relatively few novel motifs emerging in eukaryotic kinases. It is interesting to speculate why this is the case. It is possible that no new distinct mode of interaction can be accommodated at the active site or that such novel motifs are not easily reached via mutations of existing kinases. As mentioned above, kinase specificity is determined via multiple mechanisms including docking interactions, expression, localization, activation modes etc. Duplicated kinases can, therefore, be made non-redundant by diversifying the way by which they regulate their substrates to avoid miss-regulation in multiple different ways (Alexander et al. 2011). Additional research will be needed to study how the different kinase specificity mechanisms have evolved in kinases.

Protein kinases are just one of many peptide-binding domain types that can recognize diverse sets of peptide motifs. Other such domains include for example the PDZ, SH2, SH3, and WW, among many other families. It remains to be seen whether the findings described here relating to the evolution of different target motifs will apply to other such important peptide-binding domains.

## Methods

### The evolution of kinase function

Kinase domain sequences were collected for all 9 opisthokont species in *KinBase* with an annotated kinome (H. sapiens, M. musculus, *S. purpuratus, D. melanogaster, C. elegans, A. queenslandica, M. brevicollis, S. cerevisiae, C. cinerea*) (Manning et al. 2002). The kinase domain sequences were then aligned using the L-INS-i method (Katoh et al. 2005) and filtered to remove pseudokinases (kinases without expected residues at domain positions 30, 48, 123, 128, and 141). Manual corrections were then made to the multiple sequences alignment (MSA), and the *trimAl* tool employed to remove positions with 80% or more of ‘gap) characters among the sequences (Capella-Gutiérrez, Silla-Martínez, and Gabaldón 2009). Finally, a further filter was applied to remove truncated sequences with fewer than 190 kinase domain positions.

The resulting MSA (2094 sequences) was used to generate a maximum-likelihood kinase domain phylogeny with the *RaxML* tool (Stamatakis 2014). Amino acid substitutions were modelled using the LG matrix, and a gamma model was employed to account for the heterogeneity of rates between sites. A neighbour-joining phylogeny generated with the R *ape* package was used as the starting tree (Paradis and Schliep 2018).

Ancestral sequence reconstructions were performed with the CodeML program (part of the PAML package) using an LG substitution matrix (Yang 2007). No molecular clock was assumed (clock=0), and a gamma model was employed again to account for rate heterogeneity between sites. The alpha parameter of the gamma distribution was estimated (fix_alpha = 0) with a staring value of 0.5 (alpha = 0.5), and four categories of the gamma distribution were specified (ncatG=4). The physicochemical properties of the amino acids were not taken into account when performing the ancestral sequence reconstructions (aaDist = 0).

For the analysis of kinase evolution, each family and subfamily was assessed iteratively and a divergence score (*s*) was assigned to each position of the MSA. The divergence scores are calculated by comparing the family/subfamily of interest (clade A) with the closest sister clade (clade B) in the phylogeny. The score calculated is adapted from the *BADX* score of a previous publication (Edwards and Shields 2005), specifically:

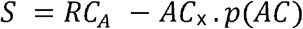

RC (recent conservation) represents the sequence conservation for the clade of interest (clade A) and is calculated here on the basis of substitution matrix similarity in the R package *bio3d (Grant et al. 2006)*. AC_X_ represents the conservation of ancestral nodes for the clade of interest (clade A) and the ancestral node for the nearest sister clade (clade B); this is given as a 1 if the predicted residues are identical to each other and a - 1 otherwise. Finally, the score is weighted by the value p(AC), which represent the the probability that the AC value was correctly assigned. For matching residues (AC=1), this is the posterior probability of the predicted residue for clade B; for differing residues (AC=-1) this is the summed posterior probability of all residues in clade B besides from the predicted residue for clade A. Therefore, scores for suspected divergence would be down-weighted if there is ambiguity concerning the nature (matching or mismatching) of the clade B ancestor.

Where the sequences of interest were divided into two or more clades in the phylogeny, only the largest clade was considered for further analysis. In some cases also the clade of interest contained spurious sequences from the wrong group, family, or subfamily. Spurious sequences were tolerated only if they comprised less than 15% of the clade sequences, otherwise the largest ‘pure) subclade (with the sequences of interest only) was selected for further analysis. For the calculation of divergence scores, the nearest sister clade to the clade of interest was selected. However, scores were only calculated if the nearest sister clade contained 5 or more sequences and belonged to the correct category (e.g. two subfamilies that are being compared must belong to the same family). All searching/manipulation of the phylogeny was performed using a custom script in R with the aid of the *ape* package.

For the global analysis represented in **Figure 2**, the number of switches was calculated at the family and subfamily level. A substitution is considered a switch if it is above the 95^th^ percentile for all subfamily (s_subfamily(95)_ =1.904) or family (s_family(95)_ = 1.793) scores. For the one-sided Fisher test)s described in the *Results* section, a site is considered to be ‘frequently switching’ if the number of switches is above the 90^th^ percentile of switch frequencies for the 246 alignment positions. This was calculated separately at the family (90^th^ percentile = 8) and subfamily (90^th^ percentile = 7) level.

### Analysis of kinases with known specificity

For the analysis of kinase specificity, 101 high-confidence specificity models of human and mouse S/T kinases were collected as described in Bradley *et al.,* 2017. Each kinase was annotated at the group, family, and subfamily level (as required) using the manual annotations given in the *kinase.com* website (Manning et al. 2002). The analysis of specificity divergence was performed separately at each of the three levels. For each level, all pairwise distances within a grouping is computed and then all possible pairwise distances are calculated between groupings. Importantly, the higher-level categorisation is retained for all pairwise comparisons. For example, at the family-level, all between-family distance comparisons would occur for kinases belonging to the same ‘group). For each pairwise comparison, the Frobenius distance between specificity models was calculated using the ‘norm() function in *R*.

### Dating the emergence of kinase groups, families, and subfamilies

First, all known kinase groups, families, and subfamilies present in animal and fungal species were retrieved from the kinase database *KinBase (Manning et al. 2002)*. The list was then filtered to remove Atypical kinases and tyrosine protein kinases. The presence or absence of each kinase group/family/subfamily across several species of the eukaryotic Tree of Life was then predicted using the *Kinannote* tool (Goldberg et al. 2013). For this purpose, we used all species from a recently published Tree of Life for which a publicly available genome/proteome sequence was available (Burki et al. 2016). The Tree of Life was then pruned in R using the *ape* package to retain these species only (55 in total). The origin of each kinase group/family/subfamily was then predicted using maximum likelihood-based ancestral state reconstruction with the *ace* function of the *ape* package (Paradis and Schliep 2018). The reported divergence times between species in the literature was then used to estimate ages for each group, family, and subfamily (Kumar et al. 2017).

Where multiple origins were predicted for a kinase group/family/subfamily, we traced the kinase emergence to the most recent common ancestral node between the predicted nodes of origin. This approach assumes no horizontal gene transfer between species or convergent evolution of kinase groups/families/subfamilies.

### Kinase motif enrichment across eukaryotic species

The phosphorylation site data was collected from a range of sources. They are as follows: *Trypanosoma Brucei* (Nett et al. 2009; Urbaniak, Martin, and Ferguson 2013), *Trypanosoma cruzi (Amorim et al. 2017; Marchini et al. 2011)*, *Leishmania infantum* (Tsigankov et al. 2013), Trichoplax adhaerens (Ringrose et al. 2013), *Homo sapiens/Mus musculus/Rattus norvegicus* (Hornbeck et al. 2015), *Strongylocentrotus purpuratus* (Guo et al. 2015), *Drosophila spp.* ((Hu et al. 2018)), *Caenorhabditis elegans (Rhoads et al. 2015)*, Magnaporthe oryzae (Franck et al. 2015), 18 fungal species (Studer et al. 2016), *Dictyostelium discoideum* (Charest et al. 2010), *Medicago truncatula* (Rose et al. 2012; Yao et al. 2014), *Glycine max* (Nguyen et al. 2012; Yao et al. 2014), *Arabidopsis thaliana* (L.-L.Lin et al. 2015; Yao et al. 2014), *Selaginella moellendorffii* (Chen et al. 2014), *Brachypodium distachyon (Lv et al. 2014)*, *Oryza sativa* (Hou et al. 2015; Yao et al. 2014), *Zea mays* (Marcon et al. 2015; Yao et al. 2014), *Chlamydomonas reinhardtii* (Wang et al. 2014), *Plasmodium falciparum/Plasmodium berghei/Toxoplasma gondii* (Invergo et al. 2017; Treeck et al. 2011), *Tetrahymena thermophila* (Tian et al. 2014), and *Phytophthora infestans* (Resjö et al. 2014).

For each species, redundant phosphosite 15mers (centred on S or T) were filtered from the analysis. Phosphorylation motifs (S/T) for each of the 48 species were obtained by running *r-motif-x* using its default parameters (p-value of *1e-06* and a minimum of 20 motif occurrences). This tool takes as its input a ‘foreground) set of known target sites and a ‘background) set of sites known not to be target sites (Wagih et al. 2016). For the background set, we randomly shuffled the flanking sequences of known phosphorylated target sites (central S/T retained). The amino acid composition of the foreground and background sets was therefore identical. This approach is expected to generate fewer spurious motif predictions than simply sampling S/T sites randomly from the proteome (Cheng et al. 2018). To generate the background set, each known target site was randomly shuffled 10 times.

For further analysis, we selected only those motifs appearing in at least a third of species within one or more superphyla (i.e. fungi, metazoa, and plants). For the excavates (three species represented here), the motif had to be present in at least two of the examined species. Motifs exclusive to the amoebozoa or heterokonts were not considered as both superphyla are represented here by only a single species. Other constraints were imposed to filter out potentially spurious motifs. Serine or threonine additions to a classical motif were not considered, as they may result from phosphosite misassignment within phosphopeptides or the clustering of phosphorylation sites in the substrate primary sequence (Moses, Hériché, and Durbin 2007; Schweiger and Linial 2010). We also considered R/K and D/E to be synonymous when identifying new motifs. Finally, D/E additions to the classic casein kinase 2 motif ‘S/T-D/E-x-D/E) were not considered as weak D/E preferences outside the +1 and +3 positions have already been described for this kinase (Sarno et al. 1997). Motifs detected here that do not match the list of motifs given in (Amanchy et al. 2007) or (Miller and Turk 2018) are declared to be ‘new’ motifs with an unknown upstream regulator.

The enrichment of kinase motifs was calculated relative to the background set of randomised peptides. The significance of motif enrichments in each species was determined by calculating binomial p-values. Here, the null probability of the motif is taken to be equal to the total frequency of motif matches (e.g. P-x-S/T-P) to the background set, divided by the total number of background matches for the superset motif (e.g. S/T-P). The calculation of equivalent frequencies for the foreground set enables an analytical p-value to be calculated using the binomial distribution. The calculated p value therefore gives an indication, for each motif, of the extent of enrichment of the motif against the background set relative to that of the most frequent superset motif (e.g. the enrichment of P-X-S.T-P relative to S/T-P).

### Number of motif matches as a percentage of the phosphoproteome

Each of the motifs previously identified using *motif-x* was screened against the known target sites of each species, and the total number target sites matching at least one motif was counted and then divided by the total number of known target sites in each species (**Supplementary Figure 1a)**. For this analysis, we do not consider motifs with only one constrained flanking position (e.g. G-S/T), where matches to the foreground set are likely to arise just by chance. These patterns may represent incomplete sequence motifs. Exceptions are made for the classic S/T-P and R-x-x-S/T signatures, which by themselves can be sufficient for kinase targeting (Errico et al. 2010; Ubersax and Ferrell 2007).

### Kinase motif enrichment for prokaryotic phosphorylation sites

The prokaryotic phosphorylation data was collected from multiple sources. Phosphorylation data for *E. coli* derives from (M.-H.Lin, Sugiyama, and Ishihama 2015; Pan et al. 2015; Potel et al. 2018). Phosphorylation data for *Sulfolobus acidocaldarius* and *Sulfolobus solfataricus* comes from the dbPSP database (Pan et al. 2015). The ‘pooled species’ in **Figure 4** represents 180 unique phosphorylation sites from 8 prokaryotic species – *Halobacterium salinarum*, *Bacillus subtilis*, *Mycobacterium tuberculosis*, *Streptomyces coelicolor*, *Escherichia coli*, *Synechococcus sp.*, *Sulfolobus solfataricus*, *Sulfolobus acidocaldarius* – all of which derives from the dbPSP database also (Pan et al. 2015).

Enrichment values and binomial p-values were calculated using the same methods described in the previous section. The *motif-x* tool was executed using its default parameters, as described above.

### Co-evolution between the kinome and phosphoproteome

A starting phylogeny for the 48 eukaryotic species was assembled using the NCBI taxonomy tool (NCBI Resource Coordinators 2018). Unresolved branches (polytomies) for particular clades were then resolved manually after referring to previous phylogenetic studies in the literature ((Cavalier-Smith et al. 2014; Drosophila 12 Genomes Consortium et al. 2007; Mathews, Tsai, and Kellogg 2000; Shen et al. 2016; Telford, Budd, and Philippe 2015)

(Cavalier-Smith et al. 2014; Drosophila 12 Genomes Consortium et al. 2007; Mathews, Tsai, and Kellogg 2000; Shen et al. 2016; Telford, Budd, and Philippe 2015)). Kinome annotations for each species were generated automatically using the *KinAnnote* tool (Goldberg et al. 2013), which employs BLAST- and HMM-based searches to identify and classify eukaryotic protein kinases.

The relationship between kinase motifs and their cognate kinases (e.g. S/T-P-x-K and CDKs) was modelled with phylogenetic independent contrasts (PIC) in R using the *ape* package (Paradis and Schliep 2018; Felsenstein 1985). This method generates phylogenetic contrasts between variables on a tree to account for the non-independence of data points (Felsenstein 1985). In **Supplementary Figure 2**, contrasts were generated for motif enrichment values on the y-axis and for relative kinase frequencies (number of kinases of interest divided by total number of kinases detected in the proteome) on the x-axis.

Tests for the phylogenetic signal of different motifs were conducted in R using the *Phylosignal* package (Keck et al. 2016). The phylogenetic plots in **Supplementary Figures 3 and 4** were also generated using *Phylosignal*.

---

**Supplementary Figure 1.**
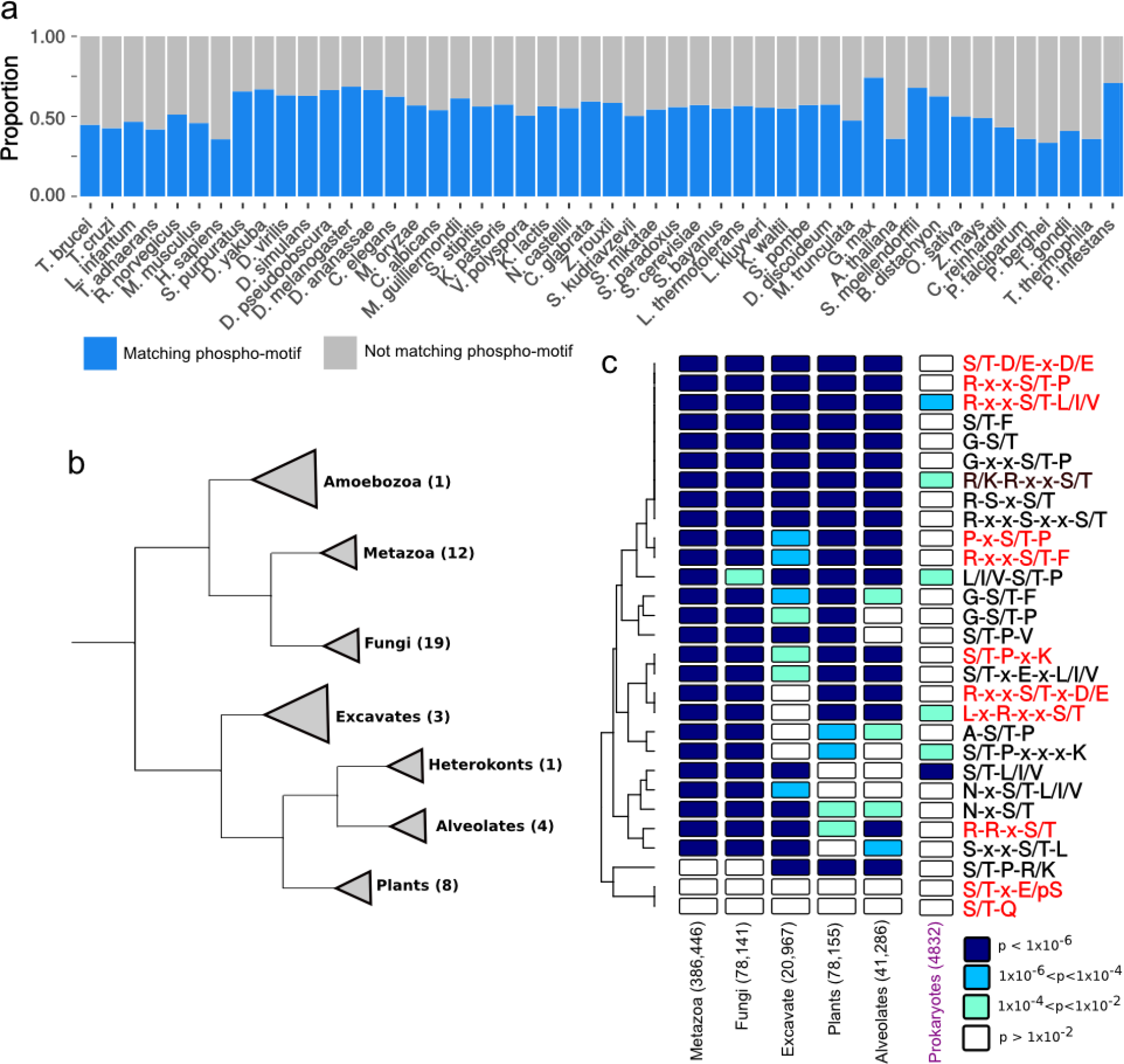
***a)*** Proportion of phosphorylation sites in each species that match a phosphorylation motif (see ***Methods***). ***b)*** A simplified version of the eukaryotic Tree of Life presented in the (Burki et al. 2016) study. The numbers in brackets correspond to the number of different species represented by phosphorylation data in this study. ***c)*** Calculation of binomial p-values (as in Figure 4) for each motif in each major clade (metazoa, fungi, plants, etc) after phosphorylation sites within a clade were pooled across species. The figure legend (bottom-right) is the same as in Figure 4.

**Supplementary Figure 2.**
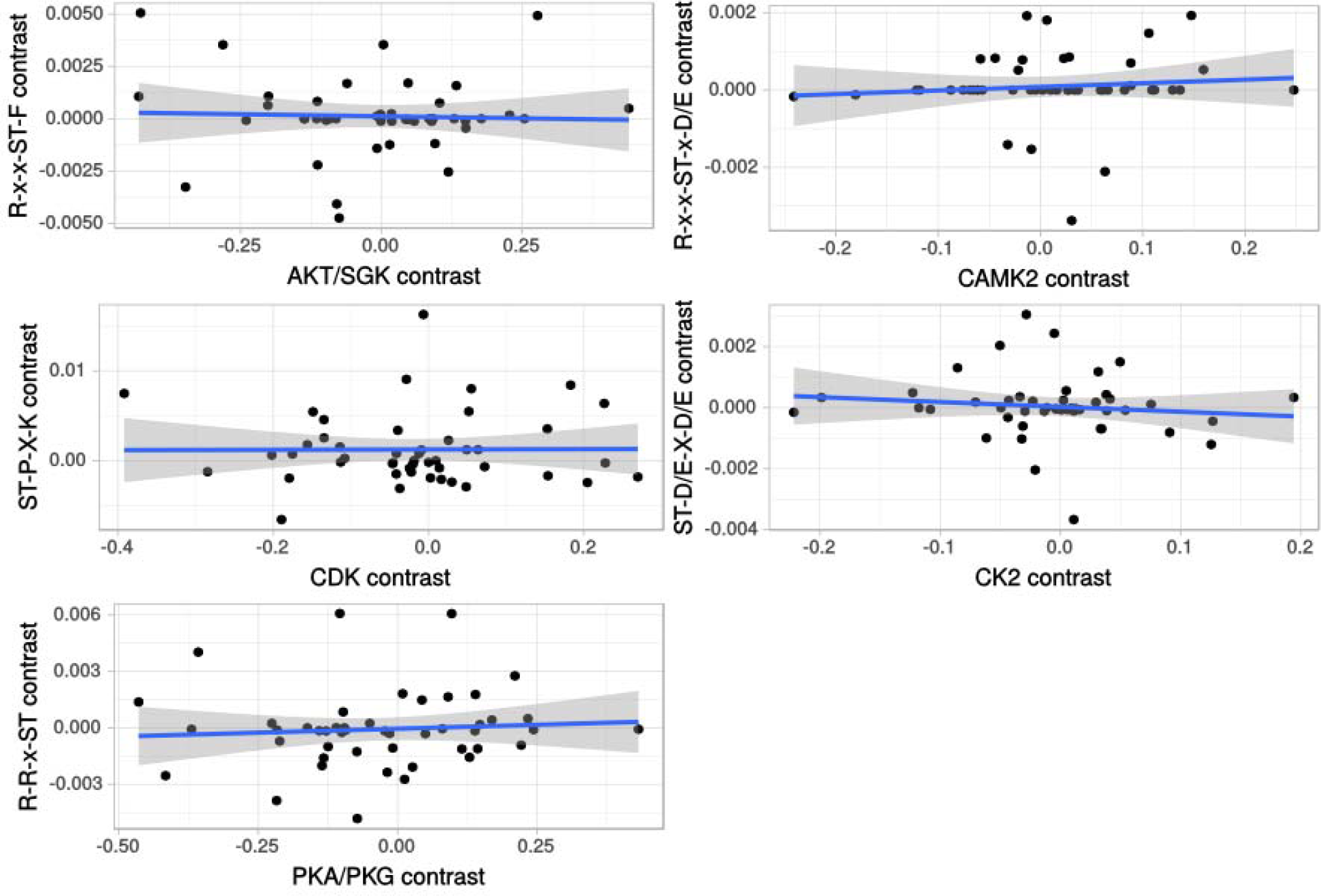
Phylogenetic independence contrasts (PIC) between five different kinase clades (AKT/SGK, CAMK2, CDK, CK2, PKA/PKG) and their corresponding substrate motifs (R-x-x-S/T-F, R-x-x-S/T-x-D/E, S/T-P-x-K, S/T-D/E-x-D/E, and R-R-x-S/T, respectively). This approach accounts for the phylogenetic non-independence between data points when comparing two continuous variables (Felsenstein 1985).

**Supplementary Figure 3.**
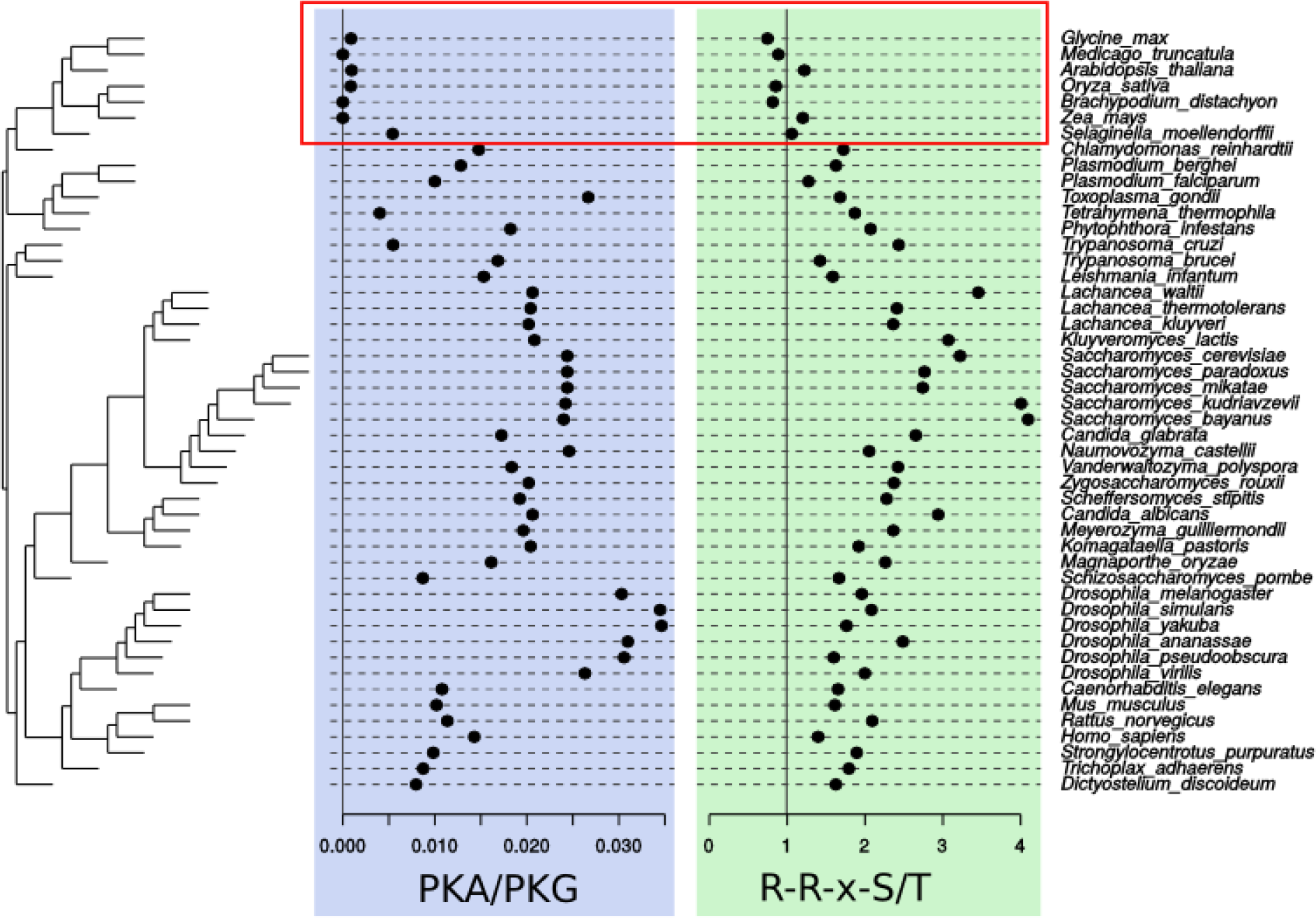
Mapping of relative kinase frequencies and substrate motif enrichments to a phylogeny of 48 eukaryotic species. Relative kinase frequencies across the 48 species were calculated for the PKA/PKG family, and motif enrichments were calculated for their cognate substrate motif (R-R-x-S/T). The red box highlights species where the absence of PKA and PKG kinases in the proteome corresponds to a lack of R-R-x-S/T motif enrichment.

**Supplementary Figure 4.**
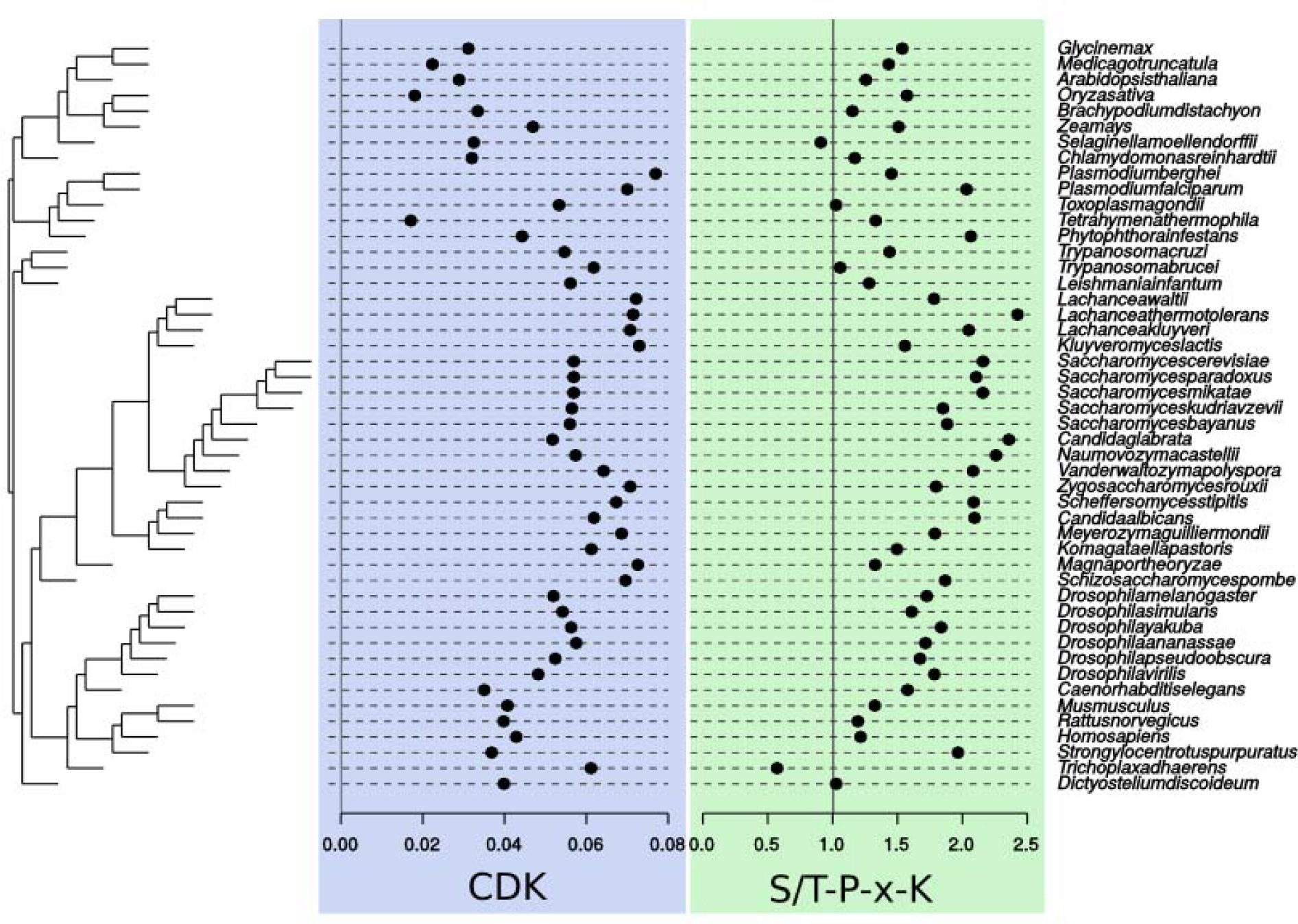
Mapping of relative kinase frequencies and substrate motif enrichments to a phylogeny of 48 eukaryotic species. Relative kinase frequencies across the 48 species were calculated for the CDK family, and motif enrichments were calculated for its cognate substrate motif (S/T-P-x-K).

**Supplementary Table 1.** Five tests for the phylogenetic signal (Cmean, I, K, K.star, and Lambda) of 9 different eukaryotic motifs. Numbers in the table represent p-values for each one of the tests. Low p-values (e.g. p < 0. 01) suggest that the motif in question is non-randomly distributed with respect to the species phylogeny of 48 eukaryotic species (as presented in Supplementary Figure 3 and Supplementary Figure 4). All tests were performed using the Phylosignal package in R (Keck et al. 2016).

|  | Cmean | I | K | K.star | Lambda |
| --- | --- | --- | --- | --- | --- |
| S/T-D/E-x-D/E | 0.062 | 0.076 | 0.175 | 0.069 | 0.001 |
| R-x-x-S/T-P | 0.086 | 0.166 | 0.067 | 0.175 | 0.001 |
| P-x-S/T-P | 0.002 | 0.003 | 0.002 | 0.002 | 0.001 |
| S/T-P-x-K | 0.001 | 0.001 | 0.001 | 0.001 | 0.001 |
| R-R-x-S/T | 0.001 | 0.001 | 0.001 | 0.001 | 0.001 |
| R-x-x-S/T-x-D/E | 0.001 | 0.001 | 0.007 | 0.001 | 0.001 |
| L-x-R-x-x-S/T | 0.001 | 0.001 | 0.002 | 0.001 | 0.001 |
| R-x-x-S/T-F | 0.006 | 0.011 | 0.052 | 0.011 | 0.001044 |
| R-x-x-S/T-L/I/V | 0.002 | 0.001 | 0.001 | 0.001 | 0.001 |

**Supplementary Table 2.** Motifs identified from phosphorylation sites in E. coli (n=2287) using the motif-x tool. In both instances, motif-x was executed using its default parameters (p<1×10^-6^ and at least 20 occurrences). The motif-x scores for each of the motifs is displayed in the second column.

| *E. coli* |  |
| --- | --- |
| Motif | Score |
| .....M[ST]..... | 15.65 |
| .....S[ST]..... | 6.18 |
| .....GK[ST]..... | 10.23 |
| ..M....[ST]..... | 4.431 |

**Supplementary Table 3.** Motifs identified from phosphorylation sites in Sulfolobus spp. (n=1655) using the motif-x tool. In both instances, motif-x was executed using its default parameters (p<1×10^-6^ and at least 20 occurrences). The motif-x scores for each of the motifs is displayed in the second column.

| *Sulfolobus spp.* |  |
| --- | --- |
| Motif | Score |
| K.....[ST]R..... | 18.37 |
| .....[ST]R..... | 8.718 |
| .....M[ST]..... | 7.955 |
| .....[ST]T..... | 5.163 |
| ...M..[ST]K..... | 9.773 |
| .....[ST].K..... | 4.584 |
| .....[ST]K..V... | 9.094 |

